# Effects of sequence motifs in the yeast 3′ untranslated region determined from massively-parallel assays of random sequences

**DOI:** 10.1101/2021.03.27.437361

**Authors:** Andrew Savinov, Benjamin M. Brandsen, Brooke E. Angell, Josh T. Cuperus, Stanley Fields

## Abstract

The 3′ untranslated region (UTR) plays critical roles in determining the level of gene expression, through effects on activities such as mRNA stability and translation. The underlying functional elements within this region have largely been identified through analyses of the limited number of native genes. To explore the effects of sequence elements when not present in biologically evolved sequence backgrounds, we analyzed hundreds of thousands of random 50-mers inserted into the 3′ UTR of a reporter gene in the yeast *Saccharomyces cerevisiae*. We determined relative protein expression levels from the fitness of a library of transformants in a growth selection. We find that the consensus 3′ UTR efficiency element significantly boosts expression, independent of sequence context; on the other hand, the consensus positioning element has only a small effect on expression. Some sequence motifs that are binding sites for Puf proteins substantially increase expression in this random library, despite these proteins generally being associated with post-transcriptional downregulation when bound to native mRNAs. Thus, the regulatory effects of 3′ UTR sequence features like the positioning element and Puf binding sites appear to be strongly dependent on their context within native genes, where they exist alongside co-evolved sequence features. Our measurements also allowed a systematic examination of the effects of point mutations within efficiency element motifs across diverse sequence backgrounds. These mutational scans reveal the relative *in vivo* importance of individual bases in the efficiency element, which likely reflects their roles in binding the Hrp1 protein involved in cleavage and polyadenylation.

## Introduction

The regulation of gene expression is central to biology, enabling functions ranging from environmental adaptation to animal development. However, deciphering the underlying logic of this regulation is difficult using only natural genetic elements because the relevant sequences in any organism vastly under-sample sequence space. For example, the roughly 6,000 genes of the yeast *Saccharomyces cerevisiae* or 20,000 human protein-coding genes are dwarfed even by the set of possible 20-mer DNA sequences (~ 1.1 x 10^12^), let alone the set of possible sequences approaching the lengths of regulatory sequences, which can span hundreds or thousands of base pairs. In addition, the regulatory sequences sampled by evolution are only a small number of the possible outcomes. Thus, additional facets of gene regulation might be learned by systematically interrogating the functional consequences of libraries of random synthetic sequences whose size vastly exceeds the number of an organism’s genes. Enabled by advances in high-throughput sequencing and oligonucleotide synthesis, this approach has been taken to develop a deeper understanding of 5′ untranslated regions (UTRs) of mRNAs [1,2], promoters [3,4] and splicing [5].

Here, we extend this massively parallel approach to the regulatory grammar of 3′ UTR sequences in the model eukaryote *S. cerevisiae*. The 3′ UTR plays important roles in mRNA metabolism, affecting mRNA stability, translation and localization [6]. These activities are mediated by proteins that bind to sequence and structural features of 3′ UTRs. Work based largely on a few well-studied yeast genes [7–9], especially *CYC1* [10–12], has identified three sequence elements in the 3′ UTR that play large roles in the determination of gene expression levels as well as 3′ end cleavage and polyadenylation. These sequence features are termed the efficiency element (consensus UAUAUA), positioning element (consensus AAWAAA, with W an A or U), and cleavage and polyadenylation site (YA_N_, with Y a C or U) [12]. Biochemical and structural investigations have shown that the efficiency element binds Hrp1 [13,14], which in turn recruits the rest of the cleavage factor I (CF I) complex. This complex is required for efficient cleavage and polyadenylation, and includes the Rna15 protein, which associates with the positioning element in the context of this complex [15,16].

Measurements of the protein levels associated with ~13,000 3′ UTR sequences, drawn largely from the yeast transcriptome along with mutant versions of 217 native sequences, demonstrated a major role for the efficiency element [17]. Studies have also interrogated yeast mRNA stabilities transcriptome-wide [18–21]. One such study [21] suggested that poly(U) elements near the 3′ end of 3′ UTRs are important determinants of stability, and hence gene expression levels, with this effect thought to be mediated by formation of RNA hairpins with the poly(A) tail. Investigations of native yeast genes have also suggested stabilizing and destabilizing roles for sequence motifs associated with binding by various RNA-binding proteins, most notably the Puf family of proteins [22–25], often via experiments involving the deletion or over-expression of Puf protein genes. In yeast, Puf proteins primarily function as repressors of gene expression via mRNA destabilization [26,27]. However, as Puf proteins act via recruitment of additional factors, Puf protein binding sites have also been found that lead to mRNA localization or increased translation [6,28,29].

As naturally-occurring 3′ UTR sequences have evolved to function in specific biological contexts, any measurement of the effects of a sequence element in a native 3’ UTR sequence context is complicated by the possible effects of co-evolved sequence features. Thus, we sought to build on the foundational studies of native yeast 3′ UTRs by performing a high-throughput assay of the expression of a single reporter gene under the regulatory control of hundreds of thousands of random 3′ UTR sequences. We determined that the efficiency element is the major regulator of gene expression, independent of sequence context. On the other hand, the positioning element and poly(U) motifs had only modest effects on expression. Three Puf protein binding site sequences were associated with substantially enhanced expression in these random sequence backgrounds, opposite to their effect in native mRNAs, pointing to a predominant role for sequence context in Puf protein-based regulation. The large number of 3′ UTR sequence variants analyzed in these experiments allowed us to determine the effects of single base pair changes in 3′ UTR elements across diverse random sequence backgrounds, indicating the relative importance of each base in efficiency element sequences.

## Results and Discussion

### Library and assay design

To assay the effects of random 50-nucleotide elements (N50) within a 3′ UTR, we generated two libraries in the context of the *HIS3* gene coding sequence and the *CYC1* gene promoter and 3′ UTR sequences. The random sequence was synthesized from equal ratios of the four nucleotides at each position and inserted into a low copy number centromeric vector. In one library (termed N50-EPC), we replaced the first 102 bases of the *CYC1* 3′ UTR with the N50 element. This element was positioned between the *HIS3* termination codon and a region of 50 bases of *CYC1* that includes the efficiency and positioning elements, the cleavage site where polyadenylation occurs, and 101 bases of constant sequence that constitute the remaining region of the *CYC1* terminator (**Fig. 1a**). In the other library (termed N50-C), the sequence 3′ of the N50 element included only the cleavage site and the same downstream constant sequence derived from the 3′ end of *CYC1* as in N50-EPC (**Fig. 1a**). Based on estimates of the number of unique transformants, the N50-EPC library consisted of 2.1 million variants and the N50-C library consisted of 2.5 million variants. Our rationale for generating these two N50 libraries was that the N50-EPC library should provide a reasonably high baseline of faithful 3′ processing through the use of the canonical *CYC1* elements, allowing the identification of random elements that would modulate gene expression around this baseline; on the other hand, the N50-C library, lacking invariant efficiency and positioning elements, was intended to have low baseline expression and thereby reveal sequence features that increase expression levels.

**Figure 1.**
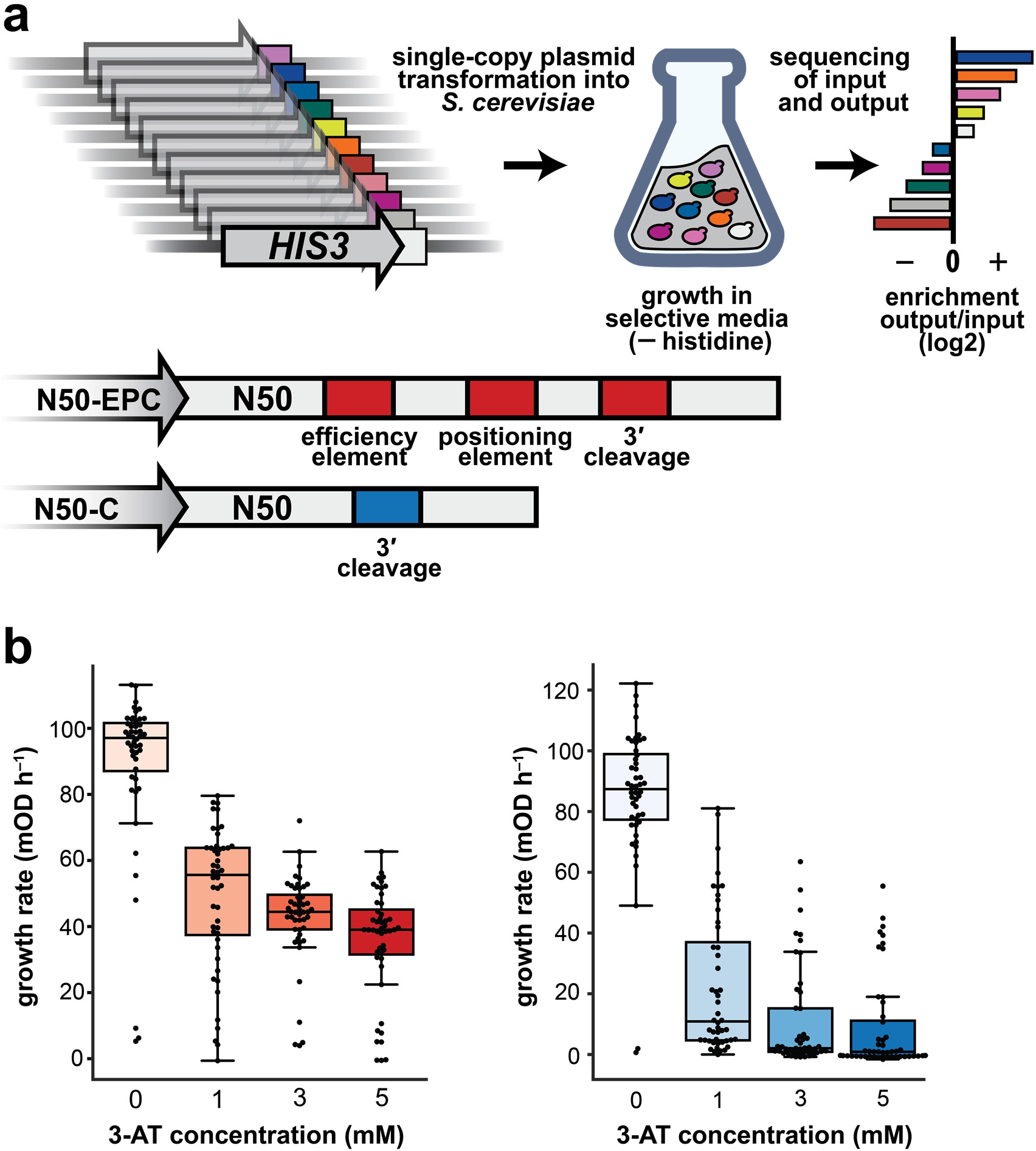
A massively-parallel assay of the effects of random 50 bp sequences on expression mediated by 3′ UTRs. (**a**) Library design and selection assay (*upper*) and library layout schematic, including known 3′ UTR motifs present in the two libraries by design (*lower*). (**b**) Determination of optimal 3-AT concentrations through growth rate measurements of 45 random variants each from the N50-EPC (*left*) and N50-C (*right*) libraries.

The plasmid-borne *HIS3* reporter gene was transformed into a yeast *his3* deletion mutant, which allowed us to read out the relative expression of library variants at the His3 protein level by selecting yeast grown in media lacking histidine and supplemented with 3-amino-1,2,4-triazole (3-AT), a competitive inhibitor of His3. By testing 45 variants from each of the two libraries over a range of 3-AT concentrations, we established that 1 mM yielded the greatest dynamic range of growth rates (**Fig. 1b**). The use of the His3 selection provided a continuous readout of protein expression that did not depend on a FACS binning strategy, as would be necessitated by fluorescence-based readouts [3,17,30,31]. The library design and selection strategy was derived from previous work to investigate 5′ UTR sequence variant effects [1] and validated in that work to report faithfully on relative His3 protein levels and growth rates of individual variants. We sequenced the N50 elements of each library prior to selection and after ~24 hr (N50-EPC) or ~30 hr (N50-C) of growth to OD_600_ = 1.0 in the absence of histidine and in the presence of 1 mM 3-AT. The relative change in abundance of each variant is presented throughout the text as a log_2_ enrichment, *Enr,* equal to log_2_(*f*_post-selection_/*f*_pre-selection_), where *f*_pre-selection_ and *f*_post-selection_ denote population frequencies of the variant before and after selection. We filtered these data to improve our confidence in the input and output variant frequencies, leaving ~590,000 N50-C sequences and ~280,000 N50-EPC sequence for which enrichment in the growth selection was quantified (see Methods).

### Overall properties of the N50-C and N50-EPC libraries

An initial analysis of sequences in the N50-C library revealed a correlation between overall AU content of the N50 element and His3 protein expression (Pearson’s *r* = 0.27; **Fig. 2a**). In contrast, the same analysis performed for the N50-EPC library showed a striking lack of correlation (Pearson’s *r* = −0.021; **Fig. 2b**). These results hinted at a greater sequence-dependence of expression in the N50-C context compared to the N50-EPC context. We thus sought to determine other 3′ UTR sequence features besides AU content that act as determinants of expression in the randomized N50 sequence backgrounds, initially by carrying out a systematic analysis of the effects of all possible 6-mer RNA sequences in the libraries (**Fig. 2c,d**). We found that the average log_2_ enrichment (*Enr*) of 6-mers ranged from 0.30 to 2.60 in the N50-C library (library mean *Enr* of 0.86), but only −0.76 to −0.39 (library mean *Enr* of −0.57) in the N50-EPC library. The range of 6-mer effects in the N50-EPC library was comparable to the uncertainties in the mean effects of each 6-mer (**Fig. 2d**, inset). Given the minimal expression consequences of N50 sequence content in the N50-EPC library, we focused our subsequent analyses on the N50-C library data.

**Figure 2.**
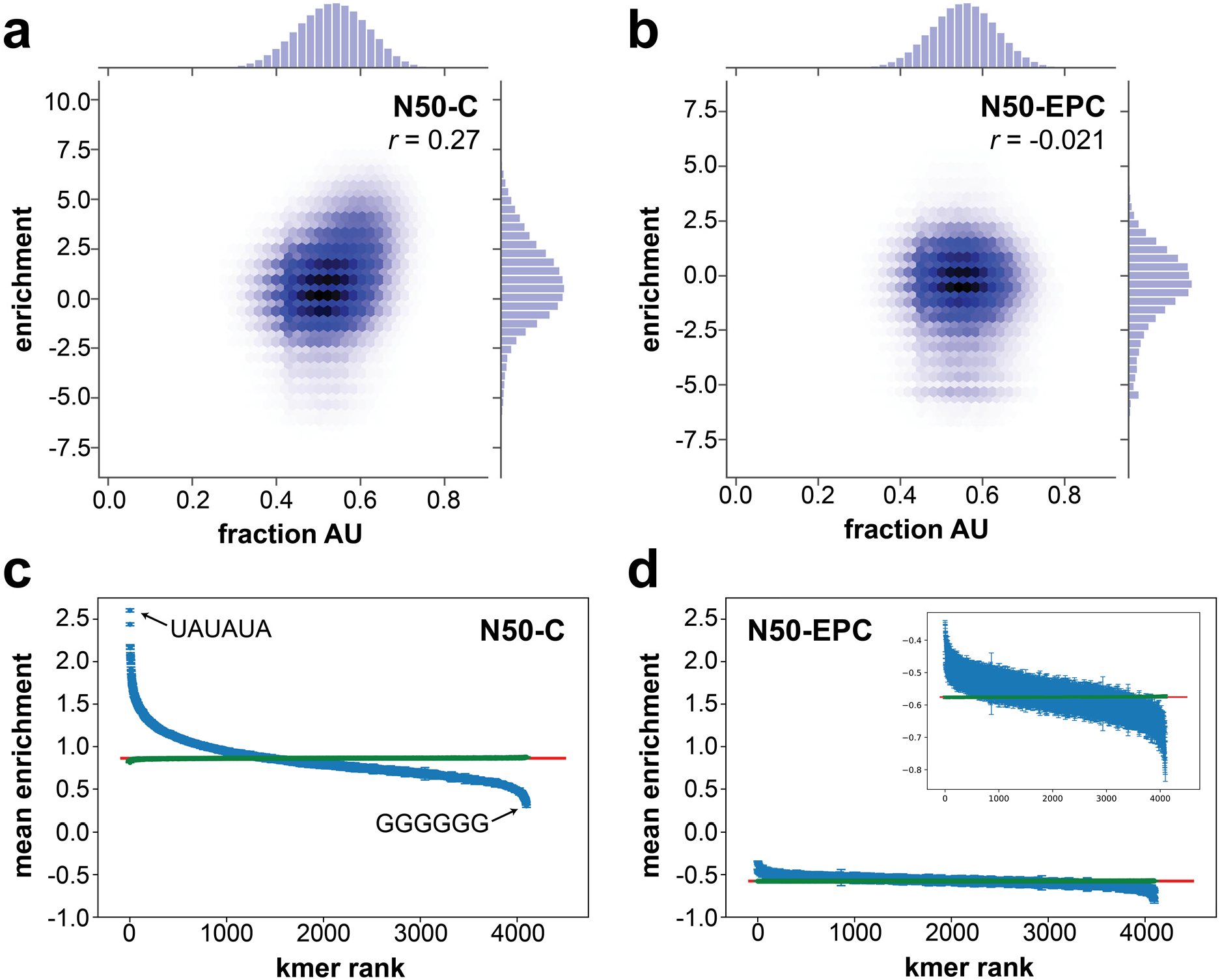
Comparison of AU content and k-mer for the N50-C and N50-EPC libraries. (**a**) and (**b**) Enrichment scores of the N50-C library (a) and the N50-EPC library (b) as a function of 50-mer sequence AU content, with values of Pearson’s *r* indicated. (**c**) and (**d**) Plots of average expression effects of all possible 6-mer sequences across the N50-C (c) and N50-EPC (d) libraries. The horizontal axis displays 6-mer sequence “rank” based on level of expression of N50 sequences containing each 6-mer (i.e., the 6-mer associated with the highest expression is assigned rank 1). Blue data, average enrichment across all library sequences containing the 6-mer; green data, average enrichment across all library sequences lacking the 6-mer; error bars, s.e.m.; red line, average enrichment across all library sequences. Inset in (d) shows 6-mer effects in the N50-EPC library on an expanded scale.

In the N50-C library, the 6-mer producing the highest average expression was UAUAUA (average *Enr* of 2.60, corresponding to an average ~6-fold enrichment in the selection across ~14,000 random sequences containing this hexamer; **Fig. 2c**). UAUAUA is the consensus efficiency element, and the next five highest-ranked 6-mer features (down to an average *Enr* of 2.09) were all point mutants of this motif. These six sequences were followed in rank by AUAUAU (*Enr* of 2.06) and the related sequence AUAUAA (*Enr* of 2.05). Lower-ranked 6-mers generally contained an increasing proportion of G and C bases. The most detrimental 6-mer was GGGGGG (average *Enr* of 0.30), with sequences such as GGAGGG, GGGAGG and GGGGGA having similar effects (average *Enr* ~0.33). These results demonstrate that the growth selection assay was capable of detecting sequence features associated with reduced, as well as enhanced, protein expression.

### Sequence determinants of efficiency element function

We sought to analyze the effects of specific motifs on relative protein expression levels, beginning with the core sequence elements previously found to be involved in cleavage and polyadenylation [12]. We first analyzed the average expression of N50 sequences carrying the canonical consensus efficiency element UAUAUA (**Fig. 3a**, middle), which as noted was associated with the largest expression boost of any 6-mer sequence (**Fig. 2c**). In contrast, shuffled sequences derived from this motif (Hamming distance of ≥3; see Methods), which have the same AU content, had a smaller effect on expression (average ~2.8-fold enrichment; **Fig. 3a**, right), demonstrating that the specific sequence of UAUAUA is necessary for maximal effect. These results confirm the generality of this motif’s importance, which had been inferred largely from native sequences and select synthetic contexts [11,12,17]. In particular, our findings demonstrate that a UAUAUA efficiency element increases protein expression regardless of sequence background, without reliance on nearby coevolved sequence features.

**Figure 3.**
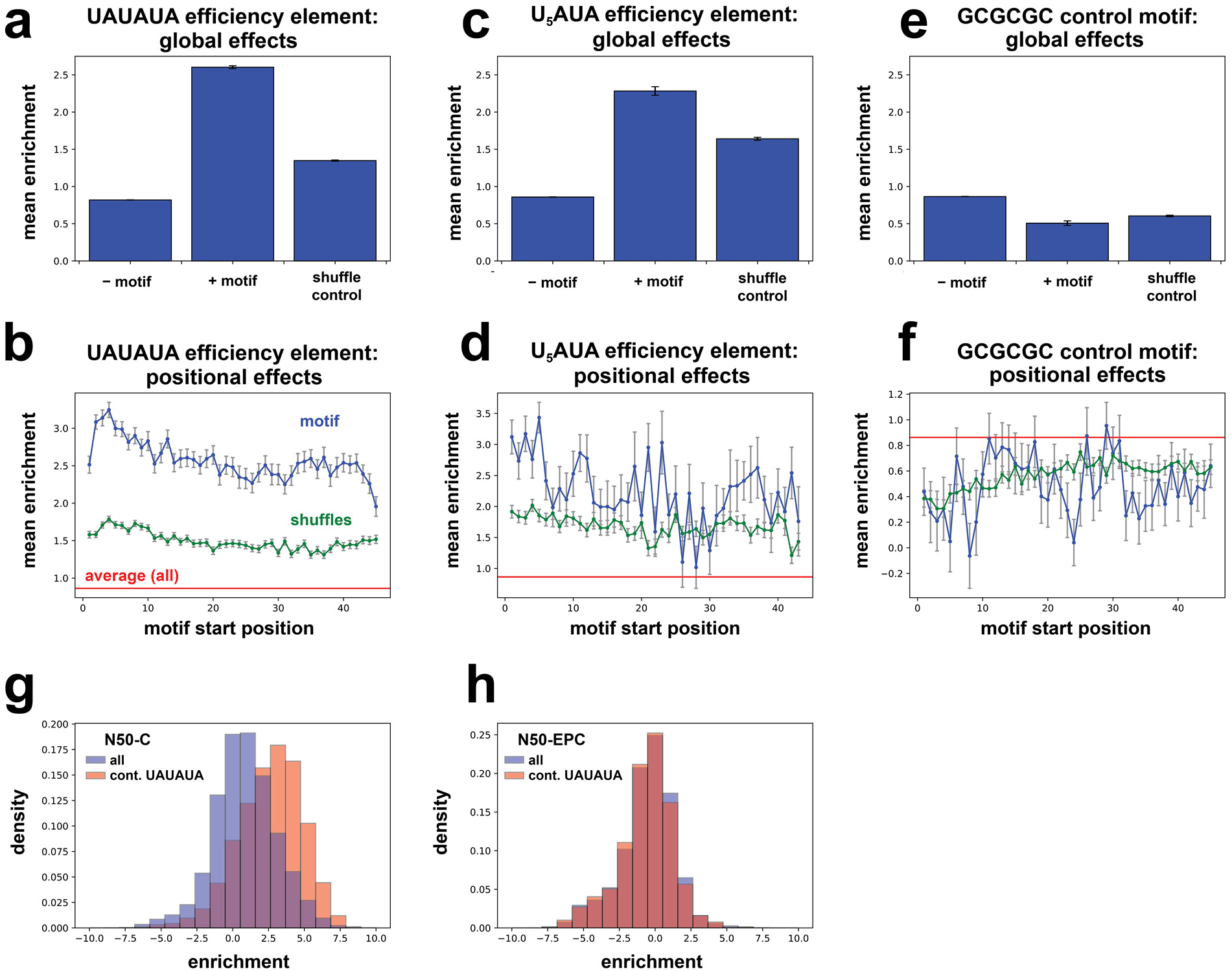
Effects of efficiency element sequences in a random context. (**a**) Average effects of UAUAUA sequence motifs on growth selection enrichment across the N50-C library. Bars correspond to (left to right): mean across all sequences lacking the indicated motif, mean across all sequences containing the indicated motif, and mean across sequences containing shuffles of the motif but not the motif itself (Hamming distance minimum = half of motif length, see Methods). (**b**) Average effects in the growth selection of UAUAUA sequence elements with 5′ end of the motif located at each position in the N50; blue, sequences containing the motif; green, sequences containing shuffles of the motif but not the motif itself (see Methods); red, average enrichment across all N50-C library sequences. (**c**) and (**d**), as for (a) and (b), respectively, but for the alternative efficiency element U_5_AUA. (**e**) and (**f**), as for (a) and (b), respectively, but for the control hexamer sequence GCGCGC. (**g**) and (**h**) Comparison of the effects of the consensus efficiency element motif UAUAUA in the N50-C library (g) and the N50-EPC library (h); enrichment score histograms of all sequences shown in blue, and of all sequences containing UAUAUA shown in tan.

The large size of the library also allowed us to determine the average effects of the consensus efficiency element when its 5′ end is located at each position in the random 50-mer. Expression mediated by the efficiency element depended on its sequence location, with N50-C variants carrying this motif generally displaying higher expression levels when the element was localized further upstream within the 3′ UTR (**Fig. 3b**, blue). However, this consensus element remained beneficial for expression at all sequence locations. The effects of shuffled hexamers derived from UAUAUA showed little position-dependence (**Fig. 3b**, green), suggesting that the shuffled sequences may largely reflect generic benefits of higher AU content (**Fig. 2a**).

To determine which effects observed with the consensus element UAUAUA generalized to other efficiency element variants, we next considered the alternative efficiency element U_5_AUA [10]. Compared to UAUAUA, U_5_AUA had similar, but weaker, expression effects, both on the average enrichment across all sequences containing this motif (*Enr* = 2.28, or ~4.9-fold, across ~1500 sequences) and as a function of location within the random 50-mer (**Fig. 3c**, middle; **Fig. 3d**, blue). Shuffled-sequence controls showed that the increase in expression associated with U_5_AUA rose above AU content effects (**Fig. 3c**, right; **Fig. 3d**, green), but to a lesser extent than for UAUAUA. As an additional control, we examined the effects of the sequence GCGCGC, the alternative pyrimidine and purine analog of the UAUAUA element. As expected, this GC-rich sequence was associated with lower-than-average enrichment (average *Enr* of 0.51, falling in the bottom 3.2% of 6-mer sequences), in a sequence context-independent and position-independent manner (**Fig. 3e,f**).

We also examined the influence of the consensus efficiency element UAUAUA on the distribution of growth selection enrichments in both the N50-C and N50-EPC libraries. Compared to the distributions across all sequences, library variants containing UAUAUA yielded a shift towards higher protein expression across the N50-C library; no such shift was observed in the N50-EPC context, which contains an efficiency element in its constant sequence (**Fig. 3g,h**). In fact, there was a small reduction in expression when an additional efficiency element was present in the N50 across the N50-EPC library (**Fig. 3h**; mean ± standard error of the mean (s.e.m.), *Enr* = −0.575 ± 0.004 across all N50-EPC sequences, vs. *Enr* = −0.648 ± 0.027 across N50-EPC sequences containing UAUAUA), suggesting that an extra efficiency element might be detrimental by reducing the efficiency of cleavage and polyadenylation. These findings suggest a “threshold model” for 3′ UTR gene regulation, in which an optimized efficiency element–positioning element–cleavage and polyA site architecture largely sets the expression level, to the exclusion of other regulatory sequences.

The selection assay results from the large 3′ UTR library effectively contained mutational scans of sequence motifs across diverse random sequence backgrounds. We sought to leverage these measurements to systematically investigate the functional role of each base in the consensus efficiency element UAUAUA. Considering first point mutations of UAUAUA that maintained AU content, we found that no such mutation yielded a larger boost in expression than the consensus sequence, as shown previously in the 6-mer analysis. Point mutations present at the 5′ and 3′ ends of the efficiency element were most detrimental to expression level compared to N50 sequences containing the unmutated consensus element, whereas the central nucleotides were the least sensitive to mutation (**Fig. 4a**). These results suggest that the most important sequence-specific binding interactions of this element with the Hrp1 protein occur at the termini. Structural work suggests that Hrp1 makes binding contacts with all six bases of the efficiency element [14], and this mutational scan informs on the relative importance and specificity of these interactions *in vivo*. Results were similar with single mutations of UAUAUA that conserve pyrimidine or purine identity instead of AU content, although a G was superior to a U at position 4 (**Fig. 4b**), with this variant being the second highest-ranked hexamer sequence (**Fig. 2c**).

**Figure 4.**
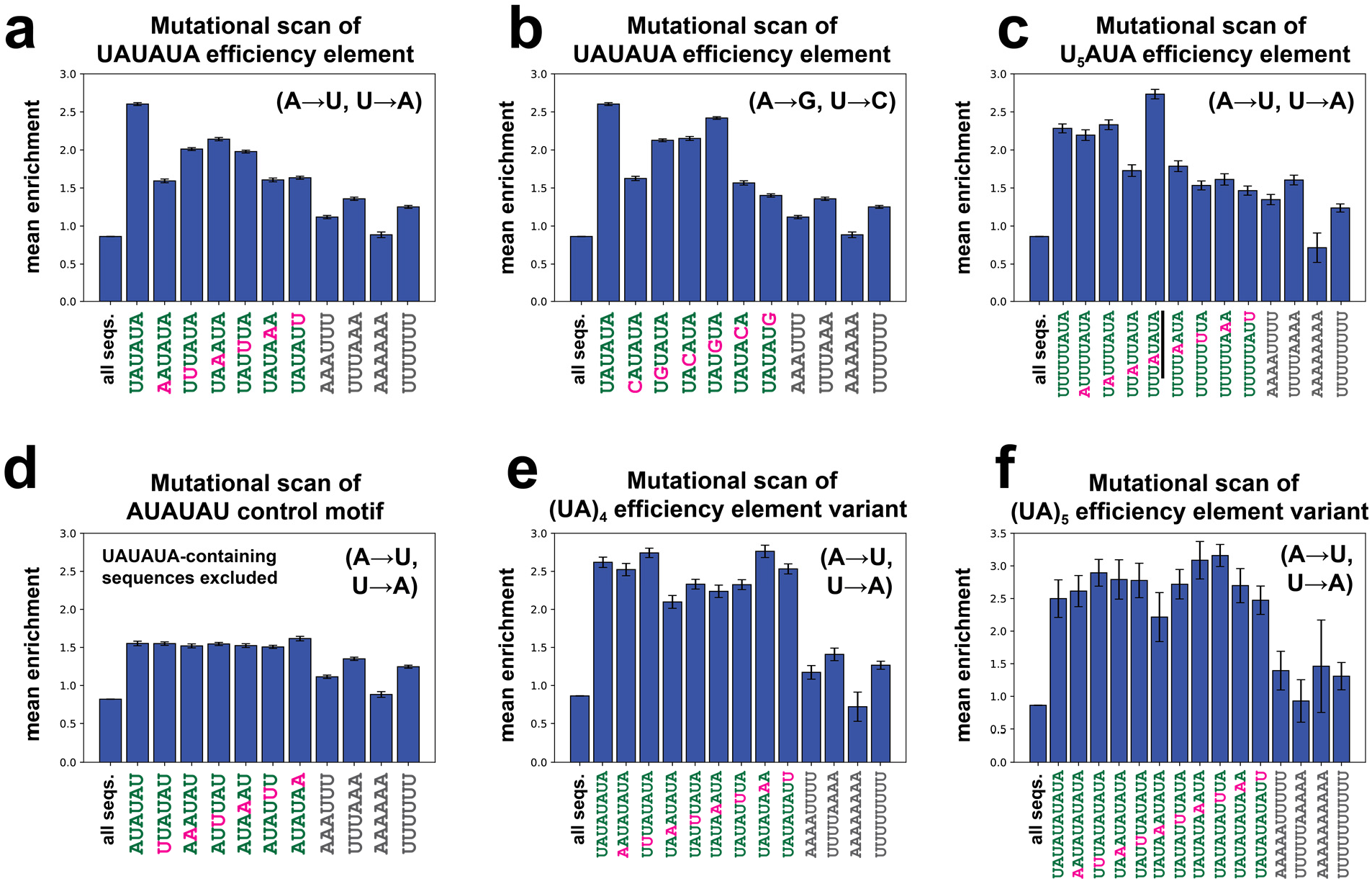
Mutational scans reveal properties of efficiency element motifs. (**a**) to (**f**) Average effects of sequence motifs (green text) and point mutants of those motifs (magenta text) on enrichment scores, compared to all sequences in the dataset (“all seqs.”) and controls with equivalent AU content (gray text). Searches for sequences containing point mutants of each motif also exclude sequences containing the consensus (unmutated) motif. (**a**) Single A→U and U→A substitutions and controls for the consensus efficiency element UAUAUA. (**b**) Single U→C or A→G substitutions and controls for the consensus efficiency element UAUAUA. (**c**) Single A→U and U→A substitutions and controls for the alternative efficiency element U_5_AUA. Underline highlights the generation of a UAUAUA efficiency element through one of the point mutants of U_5_AUA. (**d**) Single A→U and U→A substitutions and controls for the “inverted consensus” sequence AUAUAU. (**e**) Single A→U and U→A substitutions and controls for the extended efficiency element (UA)_4_. (**f**) Single A→U and U→A substitutions and controls for the efficiency element (UA)_5_.

We performed a similar analysis for the alternative efficiency element U_5_AUA. In this case, any AU content-maintaining point mutation at bases 3-8 of the motif reduced His3 expression substantially, apart from the U4A mutation that yields a consensus UAUAUA efficiency element; mutations to the first two bases had no effect (**Fig. 4c**). Furthermore, changes to bases towards the 3′ end of the motif tended to reduce expression slightly more. These findings suggest that in the case of the U_5_AUA element, the sequence U_3_AUA is in fact responsible for Hrp1 binding, consistent with the same 6-mer binding mode as observed for the consensus efficiency element. Our analysis of *k*-mer effects showed that U_3_AUA was the hexamer associated with the sixth highest expression level across the library (average *Enr* = 2.09; **Fig. 2c**), likely accounting for much of the activity in the U_5_AUA context (average *Enr* = 2.28).

A mutational scan of an AU element initiating with an A rather than a U, AUAUAU (excluding all sequences that also contain UAUAUA due to a U preceding this motif), showed that single mutations had no effect at any site (**Fig. 4d**). This result suggests that this permuted efficiency element motif may be a poor site for Hrp1 recruitment *in vivo*, despite containing a nearly-complete consensus site UAUAU, further highlighting the essential role of the nucleotides at the 5’ and 3’ termini. The insensitivity of AUAUAU to point mutations, indicating a likely lack of specificity, is striking in the context of the high expression conferred by this motif (the seventh highest-ranked hexamer, average *Enr* = 2.06).

We next considered variants of the consensus efficiency element containing additional (UA) repeats. Increasing the number of dinucleotide repeats to four or five did not further increase expression (**Fig. 4e, f**). This result provides further support for a threshold model for efficiency element function, with a single copy required for optimal expression, and saturation of this beneficial effect with the one copy. In contrast, certain (UA)_5_ mutants in which the dinucleotide repeat is broken up by point mutations did increase expression level beyond that associated with UAUAUA, with (UA)_3_U_3_A providing the highest enrichment among variants investigated (~9-fold enrichment; **Fig. 4f**). Based on these findings, (UA)_3_U_3_A may prove to be a useful generic efficiency element for achieving increased protein expression in yeast. We note that the limited increase in enrichment conferred by this motif over UAUAUA may partially be a consequence of the growth of the yeast becoming saturating under our selection conditions, as (UA)_3_U_3_A may increase expression further than the measured enrichments reflect.

### Effects of positioning element motifs and the optimal arrangement of efficiency and positioning elements

We next analyzed the positioning element, which plays a role in determining the site of cleavage and polyadenylation; mutations in this element lead to imprecise cleavage [11]. This element in yeast is A-rich, and its consensus sequence of AAWAAA bears striking similarity to the AAUAAA element found in 3′ UTRs of metazoans. Although changes in the precision by which cleavage and polyadenylation occur might be expected to affect expression by altering mRNA stability, we found that the presence of an AAWAAA element in the N50 had only modest effects on expression in the N50-C library (average *Enr* = 1.20, ~2.3-fold enrichment; **Fig 5a**, middle), not substantially higher than a hexamer of equivalent AU content, AAAUUU (**Fig 5a**, right). AAAAAA and AAUAAA, which match the AAWAAA motif, had average enrichments of *Enr* = 0.91 and 1.36, respectively. However, AAUAAA falls within the top 4% of hexamer sequences despite its modest effect size, reflecting the rapid drop in associated enrichment with hexamer rank (**Fig. 2c**). A positional analysis of the effects of AAWAAA in random sequence backgrounds showed that the positioning element generally had similar effects when found at sites throughout the N50 sequence (**Fig. 5b**).

**Figure 5.**
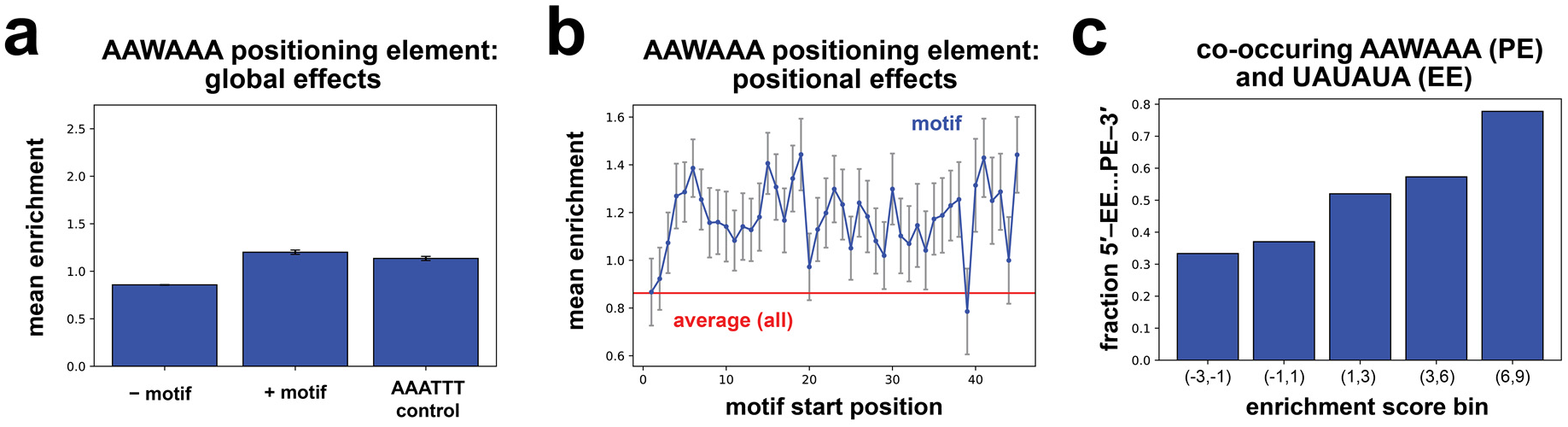
Effects of the consensus positioning element in a random sequence context. (**a**) Effects of the consensus positioning element AAWAAA on growth selection enrichment across all sequences containing the motif in the N50-C library. Bars correspond to (left to right): mean across sequences in the N50-C library lacking the indicated motif; mean across sequences containing indicated motif; and mean across sequences containing AAAUUU, a control sequence of equivalent AU content. (**b**) Average effects in the growth selection of sequences containing the AAWAAA element with 5′ end of the motif located at each position in the N50. (**c**) Among N50-C library sequences containing both a consensus efficiency element (EE) (UAUAUA) and a consensus positioning element (PE) (AAWAAA), plot of the fraction with the EE located 5′ of the PE for sequences in each enrichment score bin.

The positioning element might be expected to have strongly context-dependent effects on expression, given that binding of Rna15 protein to this element requires Hrp1 binding to a nearby efficiency element, allowing formation of the CF I complex which incorporates Rna15 [16]. To investigate the generalizability and properties of the efficiency element–positioning element interaction, we considered the fraction of N50 sequences containing the canonical consensus forms of both elements (UAUAUA and AAWAAA) in which the efficiency element is 5′ of the positioning element, across 3′ UTRs falling into different enrichment score bins. The fraction of sequences with this arrangement was larger in bins of increasingly higher enrichment scores (**Fig 5c**), suggesting that the stereotyped arrangement of these elements derived from biologically-occurring sequences is generally optimal for expression in any sequence context.

### Some Puf protein binding sites increase protein expression in a random sequence context

We examined the results of the N50-C library selection on another class of 3′ UTR sequence elements – Puf protein binding sites – including binding site motifs for Puf1 and Puf2, Puf3, Puf4, Puf5 and Puf6. By investigating shuffled versions of these motifs, we found that the Puf1 and Puf2 motif (UAAUNNNUAAU [32]) did not significantly impact His3 expression (beyond the effects of its concomitant AU content) (**Fig. 6a,b**). In contrast, Puf3 (UGUANAUA [22,29,33]) (**Fig. 6c,d**), Puf4 (UGUANANUA [22,34,35]) (**Fig. 6e,f**) and Puf5 (UGUANNNNUA [22,34]) (**Fig. 6g,h**) motifs were associated with significantly enhanced protein expression, and the Puf6 site (UUGU [36,37]) was associated with a weak increase in expression (beyond the shuffled sequence control) (**Fig. 6i,j**). The strongest increases in expression were associated with Puf motifs located closer to the 5′ end of the 3′ UTR (**Fig. 6d,f,h,j**). These findings stand in contrast to the traditional view of yeast Puf proteins as repressive elements acting mainly through mRNA destabilization (reviewed in refs. 26,27), and suggest that in the absence of additional co-evolved sequence features some Puf binding sites increase expression.

**Figure 6.**
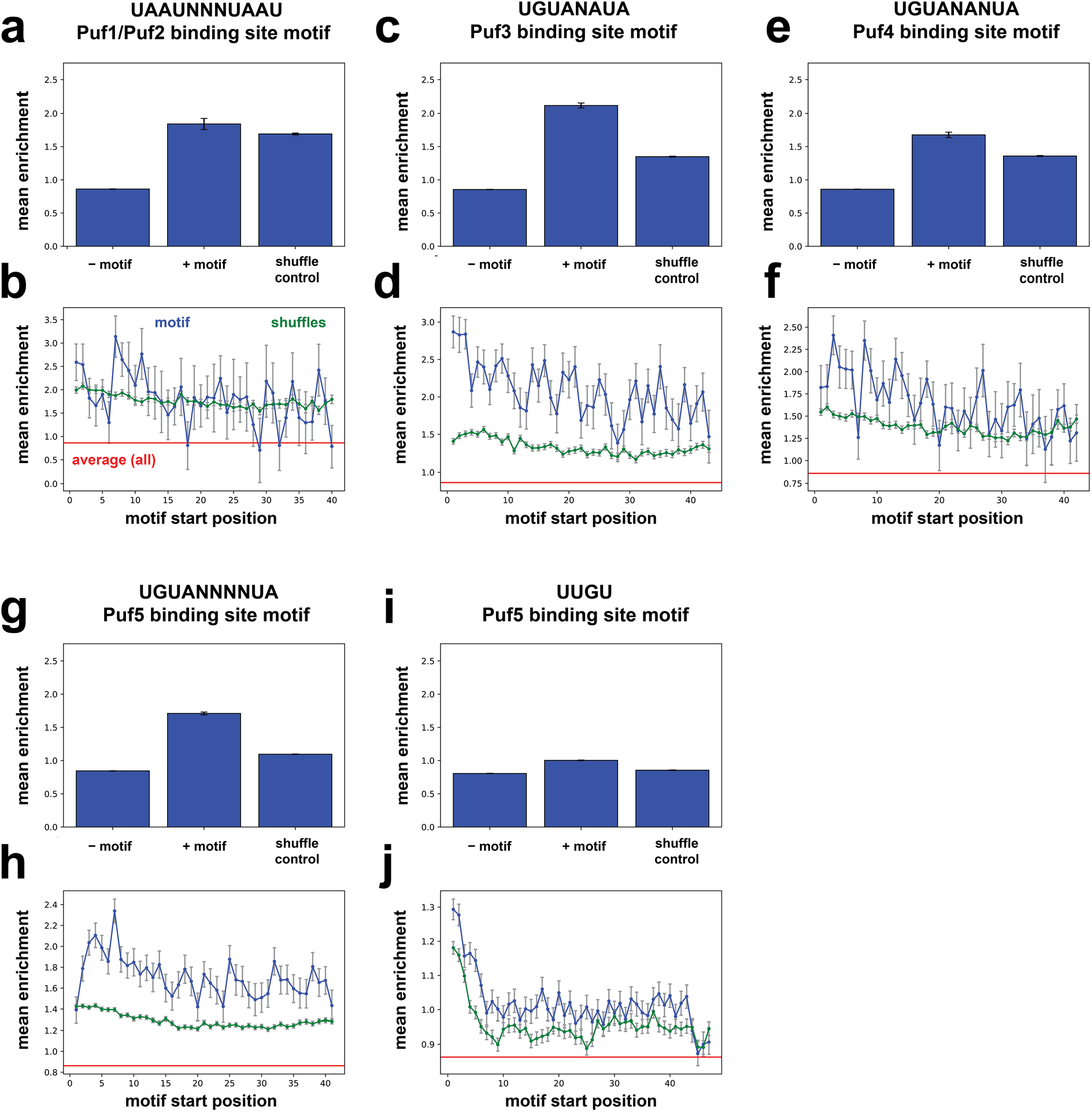
Puf protein binding sites have minimal effects when placed in a random sequence context. (**a**), (**c**), (**e**), (**g**), (**i**) Average effects of each indicated Puf binding site motif on enrichment (as in Figure 3a,c,e). Bars correspond to (left to right): mean across sequences in the N50-C library lacking the indicated motif; mean across sequences containing the indicated motif; and mean across sequences containing shuffles of the motif but not the motif itself. (**b**), (**d**), (**f**), (**h**), (**j**) Positional effects of the Puf protein binding sites indicated in (a), (c), (e), (g), and (i) above, and shuffled sequence controls, on average enrichment (as in Figure 3b,d,f). Blue, average enrichment of the motif sequence; green, average enrichment of shuffles of the motif sequence; red, average enrichment across all N50 sequences.

In the case of the Puf3 binding site, the associated enhancement of expression may be traced in part to the fact that this sequence contains UANAUA. Hence, this Puf3 site includes the consensus efficiency element UAUAUA or its point mutants at position 3, a position in this motif where mutations allowed enhancement of expression to be retained (**Fig.4a,b**). Therefore, this Puf3 binding site exemplifies a type of dual regulation based on overlapping motifs, which has been noted in 3′ UTR regulation [38,39]; in this case, the effect on expression is likely heavily influenced by strong efficiency element activity. The Puf3 binding site might thus be competed for by stabilizing and destabilizing proteins, with the relative levels of binding by Hrp1 and Puf3 likely to depend on the surrounding RNA sequence and other regulatory factors. This result may also reflect the context-dependence of the regulatory effects of Puf proteins, with Puf3 in yeast producing opposing effects – either reduced mRNA levels or increased translation – depending on metabolic state [6,29]. Biological context-dependent stabilizing or destabilizing effects have been observed with other RNA-binding proteins as well [40]. Overall, our results for the expression consequences of Puf protein motifs in the random N50 sequence background suggest that Puf protein regulation of native mRNAs depends on additional sequence context beyond the Puf binding site. These results are similar to the sequence-dependence of mRNA binding by the mouse MBNL1 and RBFOX2 proteins [41], and more broadly to other studies documenting that 3′ UTR-binding proteins associate with only a fraction of their possible binding sites *in vivo* [6,42–44]. The sequence-dependence of Puf binding site activity that we infer from our results – and the expression increase that we find was mediated by some Puf sites – may also explain why binding by the typically repressive Puf1 and Puf3 proteins is stabilizing for at least some native mRNAs [45]. Binding by Puf4 or Puf5 proteins may be generically stabilizing, or may increase levels of translation, in the absence of other sequence elements involved in recruiting destabilizing factors. We note, too, that we have not shown that the expression-boosting effect measured for Puf protein motifs was due necessarily to the binding of the cognate Puf proteins.

### Effects of poly(U) sequences on expression

A poly(U) element near the 3′ end of yeast 3′ UTRs has been implicated in stabilizing mRNA through a proposed RNA hairpin formed with the poly(A) tail [21]. In agreement with a stabilizing effect, we observed a modest average increase in His3 expression in the N50-C library for N50 sequences containing U_8_ stretches (**Fig. 7a**). However, this boost in gene expression was weaker when the U_8_ element is located in the 3′-most 25 bases of the N50 sequence (**Fig. 7a**), in contrast to the prior results [21], and instead was more substantial when the element was present in the 5′-most 25 bases (**Fig. 7a**). By calculating the average expression of sequences containing a U_8_ motif at each position in the 50-mer, we found that U_8_ increased expression most when present in the 5′ end of the N50, with weaker effects the closer the element is located to the 3′ end, and negligible effects at the 3′ terminus (**Fig. 7b**). Similar results were seen for U_6_ and U_10_ stretches (**Supp. Fig. 1**).

**Figure 7.**
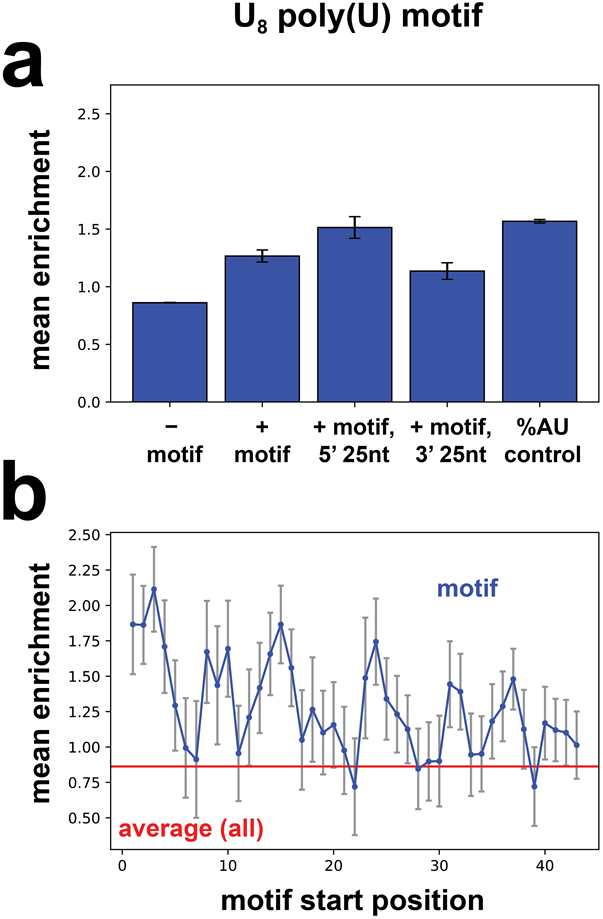
Poly(U) sequence effects on gene expression across random contexts. (**a**) Average effects of the U_8_ motif on enrichment. Plotted bars, left to right: average enrichment across sequences lacking U_8_; average enrichment across sequences containing U_8_; average enrichment across sequences containing U_8_ in the 3′-most 25 nt of the 3′ UTR; average enrichment across sequences containing U_8_ in the 5′-most 25 nt of the 3′ UTR; average enrichment of sequences containing shuffles of an 8-mer of equivalent AU content (A_4_U_4_) to compare with the effects of U_8_. (**b**) Average enrichment across all sequences containing a U_8_ motif with its 5′ end located at each N50 position (as in Figure 3b,d,f). Blue, sequences containing the motif; red, average enrichment across all N50-C library sequences.

However, the expression enhancement associated with the U_8_ sequence was smaller than the average effect of various 8-mer sequences containing 50% A and 50% U content (**Fig. 7a**), and a U_8_ sequence increased expression less than an equivalent U-rich sequence containing no more than two Us in succession **(Supp. Fig. 2**). These results make it unclear whether the protein-level effects of poly(U) sequences are specific to the mRNA-stabilizing mechanism outlined by Geisberg *et al.* [21]. These findings suggest that the documented effects of 3′ UTR sequence motifs on mRNA stability may not necessarily predict expression outcomes at the protein level, as the sequence features of 3′ UTRs influence not only RNA stability but also translation.

### Comparison of 3′ UTR sequence element effects to mRNA half-life measurements of native mRNAs

We sought to broadly compare the effects on protein expression of motifs in our random 50-mer background to the effects of these same motifs on mRNA stability in native sequence contexts. To make these comparisons, we leveraged published data on mRNA half-life across the yeast transcriptome [21], re-analyzing these data and calculating the average half-life of native yeast mRNAs containing various sequence features in their 3′ UTRs (see Methods).

We found that the increase in His3 protein expression associated with efficiency elements in the N50-C library matched an increase in native mRNA half-life, as expected (**Fig. 8a,b**), but the effect sizes were notably weaker in the native context. On the other hand, the presence of an AAWAAA positioning element sequence was associated with slightly beneficial effects on protein expression in the N50-C library, compared to slightly reduced native mRNA half-life (**Fig. 8c**). In a similar vein, GCGCGC was associated with reduced His3 protein expression, compared to an increase in native mRNA half-life (though with a large standard error; **Fig. 8d**). However, only 11 yeast genes contain a GCGCGC hexamer sequence in their 3′ UTRs, suggesting that it is evolutionarily disfavored in that context, perhaps because it typically reduces expression.

**Figure 8.**
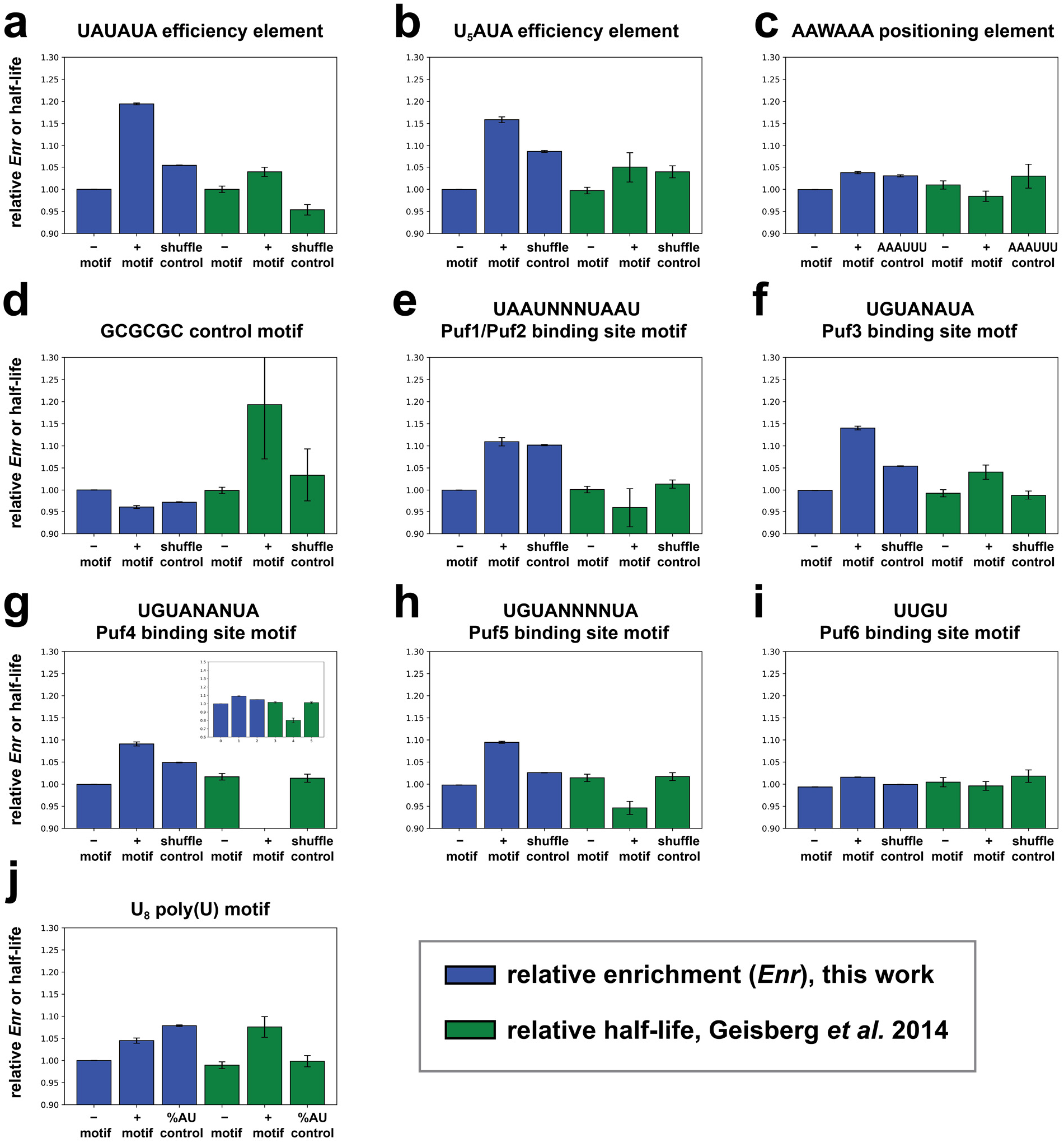
Comparison of the effects of sequence motifs on expression in a random context and on mRNA stability in native genes. (**a**) – (**j**) Average effects of the noted sequence motifs on two measurements of gene expression: relative growth selection enrichment across all sequences containing the motif in the N50-C library (blue), and relative half-lives of native yeast 3′ UTRs carrying this motif [21] (green). Triplets of bars of each color in (a) – (j) correspond to (left to right): mean across sequences lacking the indicated motif; mean across sequences containing the indicated motif; and mean across sequences containing shuffles of the motif but not the motif itself, except as noted in the following. In the case of panel (c), the third bar in each series instead represents AAAUUU, a control sequence with equivalent AU content to AAWAAA. In the case of panel (j), the third bar in each series represents average relative enrichment or relative half-life of sequences containing shuffles of an 8-mer of equivalent AU content (A4U4) to compare with the effects of U_8_.

Our analysis of the average half-lives of native mRNAs containing Puf protein binding sites showed a nominally lower half-life for the Puf1/Puf2 site (**Fig. 8e**, green), although the difference was not significant (within one s.e.m.). These findings are consistent with Puf1/Puf2 sites reducing mRNA stability on average, but with this effect subject to the wide range of native mRNA half-lives and co-evolved regulatory contexts. The effect on mRNA stability was opposite to the increase in protein expression in the random N50 context, which seems to be driven by the AU content of Puf1/Puf2 sites (**Fig. 8e**, blue). The Puf3 binding site motif was associated with a somewhat longer mRNA half-life on average, which was similar to the effects of this element on His3 protein expression, presumably reflecting the efficiency element function of this site at both the protein and the mRNA level (**Fig. 8f**). However, Cheng *et al.* [25] found that the Puf3 binding site motif UGUAAAUA was associated with reduced half-life of native mRNAs, although this same motif became stabilizing in *puf3* and *ccr4* deletion backgrounds. These results suggest that differences in mRNA half-life results between the Cheng *et al.* [25] and Geisberg *et al.* [21] studies might relate to growth conditions. The Puf4 and Puf5 binding site motifs were both associated with reduced native mRNA half-life (**Fig. 8g,h**), in contrast with the increased protein expression mediated by these elements in a random N50 context. A possible explanation for this difference is that Puf4 and Puf5 sites alone increase mRNA and protein expression levels, but additional sequence features as are present in most yeast genes result in a destabilizing effect in the native context. Alternatively, these Puf sites might affect mRNA stability and translation differentially; such differential effects might be enabled by additional roles these motifs play besides Puf protein binding. The presence of a Puf6 binding site element was associated with only a weak nominal reduction in native mRNA half-life (within one s.e.m.) (**Fig. 8i**), and a weak increase in His3 protein expression. These minimal effects may reflect the nature of Puf6 regulation, with known target genes displaying multiple Puf6 binding sites in their 3′ UTRs [36,37].

Finally, the poly(U) motif U_8_ gave strikingly different results for native mRNA half-life and His3 protein expression. Among native yeast genes the presence of a U_8_ sequence was associated with a longer half-life (**Fig. 8j**, green), consistent with Geisberg *et al.* [21], and with the analysis of U_6_A by Cheng *et al.* [25]. In contrast, as noted, U_8_ had no effect on protein expression beyond its AU content (**Fig. 8j**, blue).

### Concluding remarks

Taken together, our results indicate the importance of context in determining the expression consequences of 3′ UTR sequence features. The efficiency element emerges as a robust, context-independent regulatory sequence, with its 6-mer consensus sequence providing the largest increase in expression of any hexamer. Similarly, Puf3, Puf4 and Puf5 binding site motifs enhanced protein expression in a random context. These results suggest that an optimal efficiency element can be added to the 3′ UTR of any exogenous sequence of interest lacking this feature to increase the resultant protein expression level in yeast. Puf4 or Puf5 protein binding sites could similarly be added, although serendipitous sequence features might convert these into repressive factors; buffering the Puf motifs with surrounding random sequence might prevent this conversion. Adding AU-rich elements should also generically improve gene expression. However, the positioning element and poly(U) motifs do not display this same degree of generalizability.

As exemplified by the results for GCGCGC, the Puf binding sites and U_8_ (**Fig. 8**), the average effects of sequence elements on native mRNA stability did not generally agree with measurements of their expression effects in a random context. This lack of concordance is presumably influenced by two important factors: first, the role of evolved sequence context in modulating 3′ UTR motif function, and second, the lack of equivalence between effects on RNA level (via mRNA stability) and protein level, consistent with literature demonstrating a lack of correlation between protein and mRNA levels [46–49]. A number of factors may contribute to this regulatory complexity: multiple proteins interacting with a motif (*e.g.*, the Puf3 site; **Fig. 8f**), interactions between multiple motifs (*e.g.*, the efficiency and positioning elements, **Fig. 5c**), position-dependent effects of motifs (*e.g.*, **Fig. 3b**), and the effects of motifs in a random sequence context (*e.g.*, Puf protein binding sites, **Fig. 6**). Furthermore, sequences such as poly(U) elements and Puf protein binding motifs may affect translation in a manner distinct from mRNA stability. A dissection of the detailed interplay between these factors at both the RNA and protein levels should be a fruitful direction for efforts to decipher the underlying regulatory grammar of the 3′ untranslated region.

## Methods

### Construction of the N50-EPC and N50-C 3′ UTR libraries

We replaced the *CYC1* 3′ UTR sequence downstream of the *HIS3* stop codon on a p415-CYC1 plasmid [50] with libraries of 50-bp synthetic 3′ UTR fragments. The *CYC1* terminator is relatively short (253 bp), with well-established efficiency, positioning and cleavage sites. In the N50-EPC library, the first 102 bp were replaced with the N50 element, preserving the efficiency, positioning and cleavage elements, while in the N50-C library, the first 151 bp were replaced with the N50 element, preserving the cleavage site. The p415-CYC1-HIS3 plasmid was linearized by inverse PCR using KAPA HiFi polymerase (Kapa Biosystems) with primers F-p415-His and R-p415-HIS (oligonucleotide sequences in **Supplementary Table 1**), which remove the first 172 bp of the native *CYC1* 3’ UTR. Template DNA was digested using DpnI, and the PCR product was isolated using a DNA Clean and Concentrate Kit (Zymo Research).

The synthetic 3′ UTR fragments were constructed from Ultramer oligonucleotides (Integrated DNA Technologies) to comprise the N50-EPC or N50-C library. The oligonucleotides encoded the N50 element and either the efficiency, positioning and cleavage elements or the cleavage element. Each also encoded 20 bp of *CYC1* 3′ UTR sequence downstream of the cleavage site, as well as 30 bp of homology to the linearized backbone on both the 5′ and 3′ ends, for cloning by Gibson assembly [51]. The oligonucleotides were used as PCR templates and amplified by six rounds of PCR using KAPA HiFi polymerase (Kapa Biosystems) and primers F_N50_lib and R_N50_lib. We limited the cycles of PCR amplification to maintain sequence diversity in the libraries. After amplification, the PCR product was isolated using a DNA Clean and Concentrate Kit (Zymo Research).

The final libraries were assembled using Gibson assembly [51]. Briefly, four 20 μL reactions each containing 100 fmol of plasmid backbone, 200 fmol of 3’ UTR library and 10 μL of NEB HiFi Builder 2x master mix were incubated at 50 °C for 1 hour. Reactions were pooled and isolated using a DNA Clean and Concentrate Kit (Zymo Research), and samples were used to transform by electroporation 40 μL of Electromax DH10B *E. coli* (Agilent). Dilutions of 1:1000 and 1:10,000 were plated on LB agar plates supplemented with 100 μg/mL ampicillin to estimate the number of unique transformants in each library. The N50-EPC library contained approximately 4 x10^6^ transformants, and the N50-C library contained approximately 3.4 × 10^6^ transformants. The remaining cells transformed with library were shaken overnight at 37 °C in LB media supplemented with 100 μg/mL ampicillin, and the plasmid library was isolated using a miniprep kit (Qiagen).

### Yeast transformation

The N50-EPC and N50-C libraries were transformed into BY4741 *his3*::*KanMX* using a high-efficiency yeast transformation protocol [52]. Briefly, a 5 mL of culture was grown overnight at 30 °C in YEPD. The saturated culture was back-diluted into 50 mL of fresh 2x YEPD to an approximate OD_660_ of 0.1. Cultures were grown at 30 °C for approximately six hours, until the OD_660_ reached approximately 1.0. Cells were pelleted, resuspended in 10 mL of water, split into ten separate microcentrifuge tubes and pelleted again. Cells in each tube were resuspended in 36 μL of 1M LiAC, 240 μL of 50% w/v PEG 3350, 50 μL of 2 mg/mL salmon sperm carrier DNA that had been denatured by boiling and 200 ng of plasmid miniprep in 36 μL of water. Tubes were transferred to a 42 °C water bath and incubated for 40 min. Cells were pelleted, resuspended in 1 mL of water, combined into a single tube, and dilutions of 1:1,000 and 1:10,000 were plated on SD-Leu agar plates and grown 48 h at 30 °C to estimate the number of unique transformants. The remaining cells were diluted into 200 mL of SD-Leu media and grown overnight with shaking at 30 °C. Aliquots of 10 mL of culture were pelleted, resuspended in 1 mL of SD-Leu, mixed with 300 μL of 50% glycerol and stored at −80 °C.

### Growth curve experiments

For each of the N50-EPC and N50-C libraries, 45 random colonies transformed with the library and three colonies transformed with a reporter plasmid with the *CYC1* terminator were used to inoculate 200 μL of SD-Leu media in a 96-well plate. The colonies were shaken overnight at 30 °C in a Biotek Synergy H1 plate reader. Two μL of each saturated culture was used to inoculate 200 μL of SD-Leu -His media supplemented 0, 1, 3, or 5 mM 3-AT and shaken for 48 h at 30 °C in a Biotek Synergy H1 plate reader, with OD_660_ measured every 15 min. The maximum growth rate for each random library member was determined by calculating the most rapid increase in OD_660_.

### Massively-parallel growth selection assay for His3 expression

Glycerol stocks of each library stored at −80°C were thawed and used to inoculate 100 mL of SD-Leu media. Cultures were grown overnight at 30 °C, and 5 mL of each culture was stored at 4 °C to serve as the input sample for the selection. The OD_660_ of each library was measured and approximately 2 ×10^8^ cells were used to inoculate 100 mL of SD-Leu-His media supplemented with 1 mM 3-AT. Each culture was shaken at 30 °C until the OD_660_ measured approximately 1.0. (~24 hours for the N50-EPC library and ~30 hours for the N50-C library). 5 mL of post-selection culture was stored, and plasmids from both before and after selection were isolated using the Yeast Plasmid Miniprep II Kit (Zymo Research).

### Preparation of sequencing libraries

Sequencing libraries were prepared as 225 bp amplicons containing the 3′ UTR libraries. Plasmids isolated before and after selection were amplified by 12-16 cycles of PCR using primers that contained Illumina adapter sequences and unique sequencing indices. PCR products were isolated using a DNA Clean and Concentrate Kit (Zymo Research) and quantified using a Qubit fluorometer. The sequencing libraries were diluted to 2 nM and denatured for sequencing following the standard Illumina protocol. DNA sequencing was performed on an Illumina Nextseq 550 instrument sequencer. To identify the set of sequences in our library, we made use of the program Bartender, which collapses similar sequences into a set of consensus sequences [53]. We ran Bartender using the following options: -t 40 -d 8 -z −1 -c 1 -l 8. This set of consensus sequences was used in all subsequent analyses, with alignments performed to these sequences using Bowtie2 [54].

### Analysis of the effects of sequences in the 3′ UTR on gene expression in the N50-C and N50-EPC libraries

For random 6-mer sequences, a custom script was used to generate a list of all possible hexamer RNA sequences and then to determine the mean enrichment (and its standard error) for the subset of library sequences containing each hexamer in the N50 sequence. The same calculations were performed for the subset of library sequences lacking each hexamer in the N50 sequence. Such calculations were performed both for the N50-C and the N50-EPC library. The resulting lists of 6-mer sequences and associated average enrichments were then sorted by enrichment of sequences containing each 6-mer to determine hexamer ranking in each library.

For known 3′ UTR elements, the average effects of specific sequence elements on growth selection enrichment were calculated by using a custom script to determine mean enrichment of the subset of library sequences containing the sequence element(s) in question in the N50 sequence, making use of a string search for each element across the N50 sequences in the library. The average of sequence elements located at a specific position in the N50 were calculated as follows. FIMO [55] was used to determine the locations of all instances of a perfect match to the motif of interest in the library. Locations of each shuffled form of each motif of interest were determined in the same manner. FIMO was run with a uniform background and a *p*-value threshold set at just above the expected probability of the motif in question emerging at random (*e.g.*, (0.25)^6^ for UAUAUA). The positionally segregated average effects of each motif were determined by using a custom script to determine mean enrichment of the subset of library sequences containing the sequence element in question with the motif 5′ end (“start” sequence output from FIMO) located at each position in the N50 sequence. These analyses were also performed with shuffled sequences derived from motifs of interest. Shuffled sequences were generated using a custom Python script. Output shuffled sequences were filtered for the criterion that they be a Hamming distance of at least half the motif length away from the starting sequence (e.g., Hamming distance of 3 for the motif UAUAUA) unless otherwise noted. The number of shuffles considered for each sequence element was as many as possible matching the above criteria, up to a maximum of 50 shuffles, unless otherwise noted.

### Analysis of mRNA half-life effects of sequence elements in native yeast gene 3′ UTRs

Several existing datasets describe mRNA stability across native genes. We chose to compare our relative protein expression data from the growth selection experiments to mRNA half-life data generated with a direct RNA sequencing approach [21]. We matched the mRNA sequence from the S288C reference genome with its corresponding isoform using the gene name and 3′ UTR length with a custom Python script. Because the relative abundance of each isoform detected is not reported, we used the 3′ UTR isoform with half-life nearest to the reported mean half-life of the gene as the representative 3’ UTR sequence to avoid including low abundance transcripts in our analyses. This procedure resulted in a list of 3547 representative 3′ UTR isoforms (one per gene) and their associated half-lives.

### Comparison of sequence motif effects on native gene mRNA half-life vs. random library protein expression

We compared the consequences of a number of sequence elements on relative protein expression (growth selection enrichment score) in the N50-C 3′ UTR library to the consequences of these same elements on mRNA half-life across native 3′ UTRs in *S. cerevisiae*, based on the dataset [21] described in the previous section. Relative half-lives associated with each 3′ UTR (and the associated mRNAs) were calculated as (λ(3′ UTR) – < λ >) / < λ >, where λ denotes half-life and < λ > denotes the average half-life across all genes in the data set. Similarly, relative enrichments were calculated as (*Enr*_(norm)_ – <*Enr*_(norm)_>) / < *Enr*_(norm)_ >, where *Enr*_(norm)_ is the normalized enrichment in the growth selection and < *Enr*_(norm)_ > is the average normalized enrichment across all sequences in the N50-C library. The normalized enrichment *Enr*_(norm)_ was calculated as *Enr* – *Enr*_(min)_, where *Enr*_(min)_ is the lowest enrichment among all sequences in the library. Normalized enrichment was used in the relative enrichment calculations to produce a quantity that is always positive.

Average effects of specific sequence elements on relative mRNA half-life were calculated by using a custom Python script to determine mean relative half-life of the subset of native gene 3′ UTR sequences containing the sequence element(s) in question, using a string search of the UTR sequences for each motif of interest. Similarly, the average effects of specific sequence elements on relative enrichment in the growth selection were calculated by using the same custom script to determine mean relative enrichment of the subset of N50-C or N50-EPC library 3′ UTR sequences containing the sequence element(s) in question.

## Authors’ Contributions

B.M.B., B.E.A., J.C., and A.S. performed experiments. A.S., B.M.B., J.C., and B.E.A. analyzed the data. A.S. and S.F. wrote the manuscript with input from B.M.B., J.C., and B.E.A.

## Acknowledgements

We thank members of the Fields lab for helpful discussions.

## Supplementary Figures

**Supplementary Figure 1.**
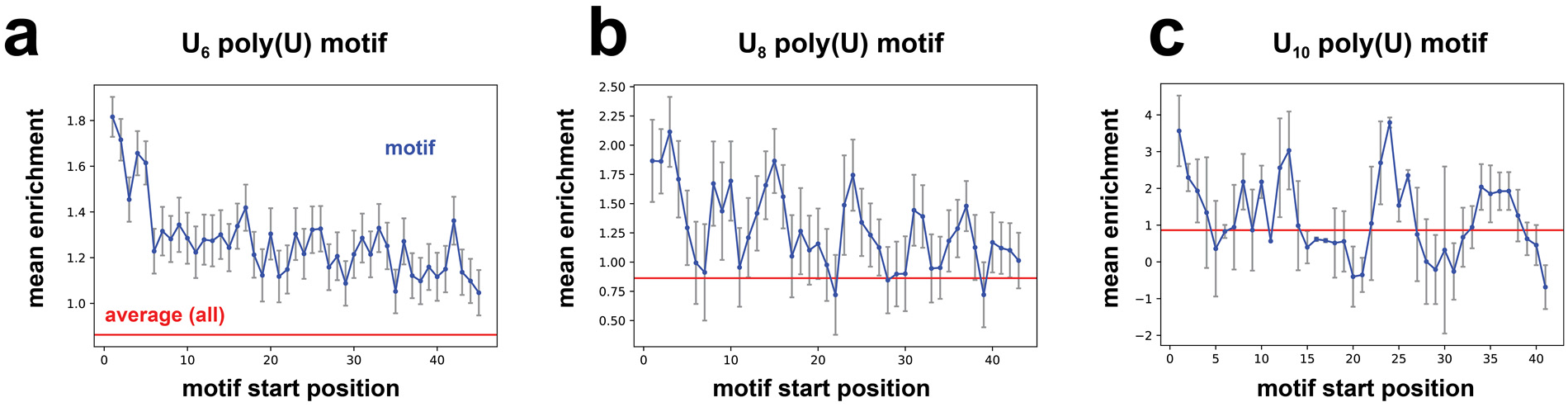
Average per-position growth selection enrichment (as in Figure 3b,d,f) of N50C 3′ UTR library variants containing poly(U) motifs with 6 (a), 8 (b), or 10 (c) sequential U nucleotides. Blue, sequences containing the motif; red, average enrichment across all N50-C library sequences.

**Supplementary Figure 2.**
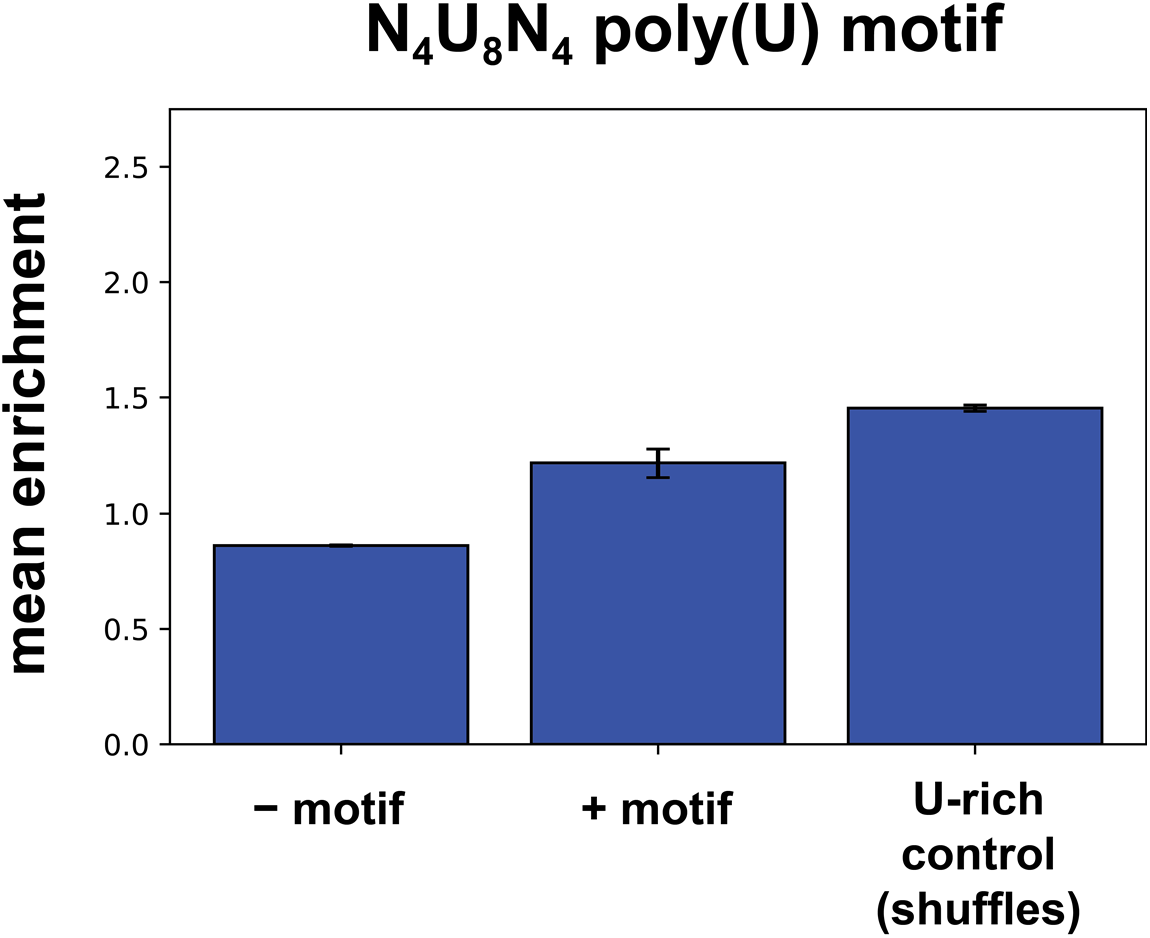
Comparing the gene expression effects of poly(U) sequences with those of equally U-rich sequences. Bars correspond to (left to right): mean enrichment across sequences in the N50-C library lacking N_4_U_8_N_4_; mean enrichment across sequences containing N_4_U_8_N_4_; and mean enrichment across sequences containing shuffles of N_4_U_8_N_4_ with no more than two sequential U’s but not N_4_U_8_N_4_ itself, while also excluding all sequences containing the consensus efficiency element UAUAUA.

## Supplementary Tables

**Supplementary Table 1:**
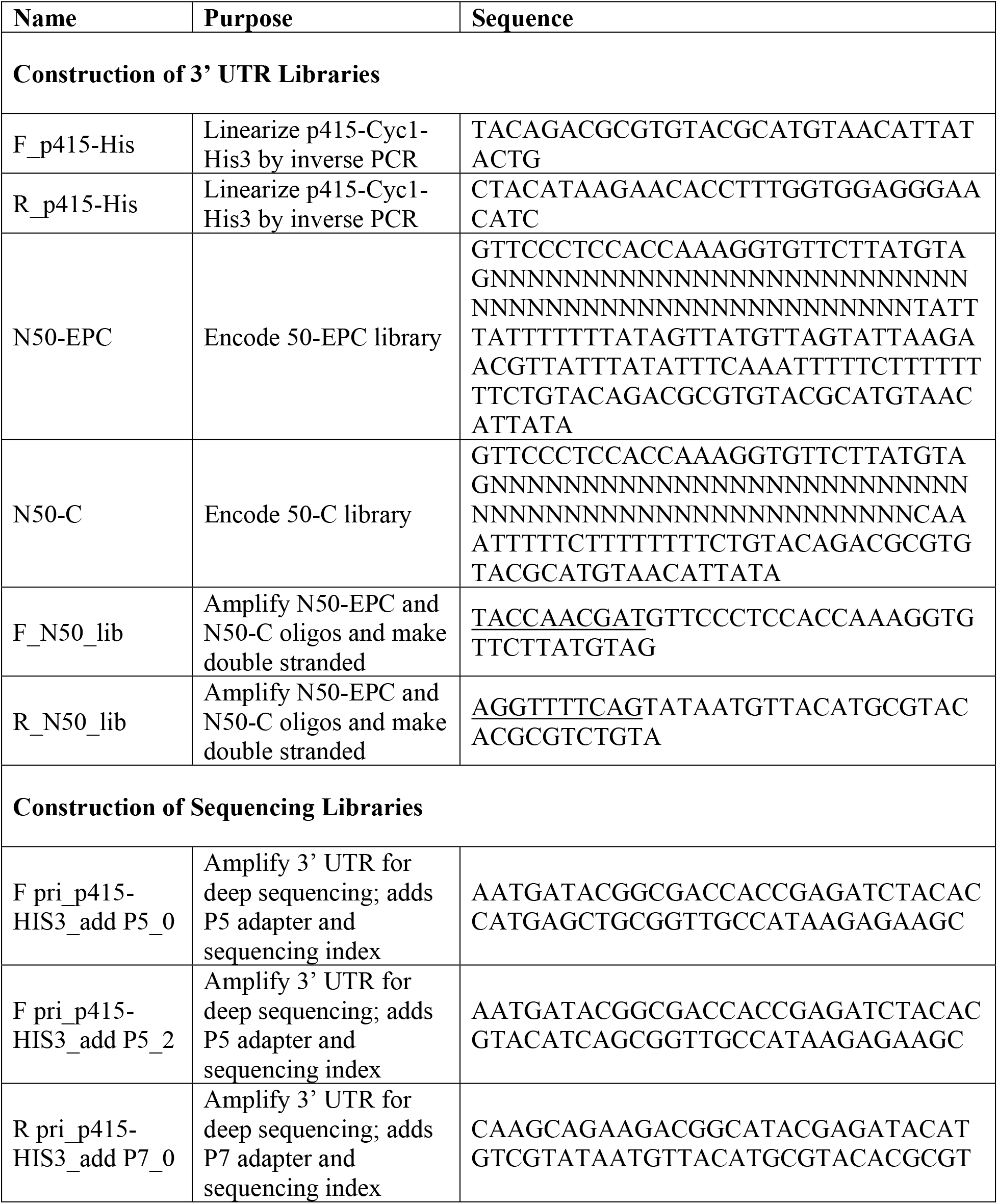
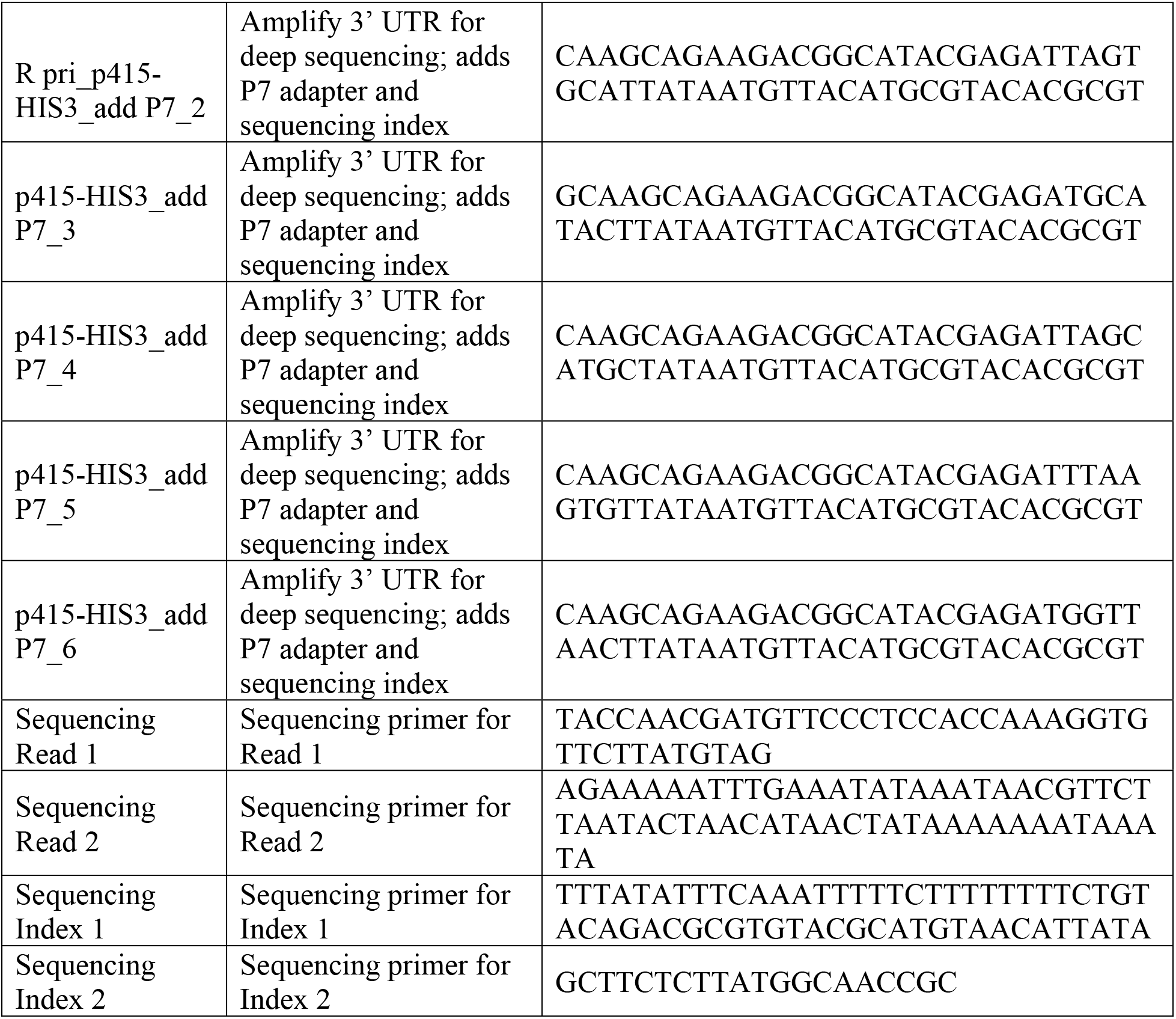
Oligonucleotide sequences.

